# An Algorithm for Cellular Reprogramming

**DOI:** 10.1101/162974

**Authors:** Scott Ronquist, Geoff Patterson, Markus Brown, Stephen Lindsly, Haiming Chen, Lindsey A. Muir, Max Wicha, Anthony Bloch, Roger Brockett, Indika Rajapakse

## Abstract

The day we understand the time evolution of subcellular elements at a level of detail comparable to physical systems governed by Newton’s laws of motion seems far away. Even so, quantitative approaches to cellular dynamics add to our understanding of cell biology, providing data-guided frameworks that allow us to develop better predictions about, and methods for, control over specific biological processes and system-wide cell behavior. In this paper, we describe an approach to optimizing the use of transcription factors (TFs) in the context of cellular reprogramming. We construct an approximate model for the natural evolution of a cell cycle synchronized population of human fibroblasts, based on data obtained by sampling the expression of 22,083 genes at several time points along the cell cycle. In order to arrive at a model of moderate complexity, we cluster gene expression based on the division of the genome into topologically associating domains (TADs) and then model the dynamics of the TAD expression levels. Based on this dynamical model and known bioinformatics, such as transcription factor binding sites (TFBS) and functions, we develop a methodology for identifying the top transcription factor candidates for a specific cellular reprogramming task. The approach used is based on a device commonly used in optimal control. Our data-guided methodology identifies a number of transcription factors previously validated for reprogramming and/or natural differentiation. Our findings highlight the immense potential of dynamical models, mathematics, and data-guided methodologies for improving strategies for control over biological processes.

**Significance Statement:** Reprogramming the human genome toward any desirable state is within reach; application of select transcription factors drives cell types toward different lineages in many settings. We introduce the concept of data-guided control in building a universal algorithm for directly reprogramming any human cell type into any other type. Our algorithm is based on time series genome transcription and architecture data and known regulatory activities of transcription factors, with natural dimension reduction using genome architectural features. Our algorithm predicts known reprogramming factors, top candidates for new settings, and ideal timing for application of transcription factors. This framework can be used to develop strategies for tissue regeneration, cancer cell reprogramming, and control of dynamical systems beyond cell biology.

## Introduction

In 1989, pioneering work by Weintraub *et al.* successfully reprogrammed human fibroblast cells to muscle cells via over-expression of transcription factor (TF) MYOD1, becoming the first to demonstrate that the natural course of cell development could be altered [1]. In 2007, Yamanaka *et al.* changed the paradigm further by successfully reprogramming human fibroblast cells to an embryonic-stem-cell-like state (induced pluripotent stem cells; iPSCs) using four TFs: POU5F1, SOX2, KLF4, MYC. This showed that a differentiated cell state could be reverted to a more pluripotent state [2].

These remarkable findings demonstrated that the genome is a system capable of being controlled via an external input of TFs. In this context, determining how to push the cell from one state to another is, at least conceptually, a classical problem of control theory [3]. The difficulty arises in the fact that the dynamics – and even proper representations of the cell state and inputs are not well-defined in the context of cellular reprogramming. Nevertheless, it seems natural to treat reprogramming as a problem in control theory, with the final state being the desired reprogrammed cell. In this paper, we provide a control theoretic framework based on empirical data and demonstrate the potential of this framework to provide novel insights into cellular reprogramming [4, 5].

Our goal is to mathematically identify TFs that can directly reprogram human fibroblasts to a desired target cell type. As part of our methodology, we create a model for the natural dynamics of proliferating human fibroblasts. We couple data from bioinformatics with methods of mathematical control theory–a framework which we dub *data-guided control* (DGC). Using time series data and a natural dimension reduction through topologically associating domains (TADs), we capture the natural dynamics of the cell, including the cell cycle.

We use this model to determine a principled way to identify the best TFs for efficient reprogramming of a given cell type toward a desired target cell type. Previously, selection of TFs for reprogramming has been based largely on trial and error, typically relying on TF differential expression between cell types for initial predictions. Recently, Rackham *et al.* devised a predictive method based on differential expression as well as gene and protein network data [6]. Our approach is fundamentally different in that we take a dynamical systems point of view, opening avenues for investigating efficiency (probability of conversion), timing (when to introduce TFs), and optimality (minimizing the number of TFs and amount of input).

Using genomic transcription and architecture data, our method identifies TFs previously found to reprogram human fibroblasts into embryonic stem cell-like cells and reprogram fibroblasts into muscle cells. Our method also predicts TFs for conversion between human fibroblasts and many additional target cell types. In addition, we show the efficacy of using TADs for genome dimension reduction. Our analysis predicts the points in the cell cycle at which the insertion of TFs can most efficiently affect a desired change of cell state. Implicit in this approach is the notion of distance between cell types, which is measured in terms of the amount of transcriptional change required to transform one cell type into another. In this way, we are able to provide a comprehensive quantitative view of human cell types based on the respective distances between cell types. Our framework separates into three parts:

1. **Define the state.** Use structure and function observations of the initial and target cell types’ genomes to define a comprehensive state representation.
2. **Model the dynamics.** Apply model identification methods to approximate the natural dynamics of the genome, from time series data.
3. **Define and evaluate the inputs.** Infer from bioinformatics (TF binding location and function) where TFs can influence the genome, then quantify controllability properties with respect to the target cell type.

The actual dynamics of the genome are undoubtedly very complicated, but as is often done in mathematical modeling studies, we use measurements to identify a linear approximation. This will take the form of a difference equation that is widely studied in the control systems literature, [7]

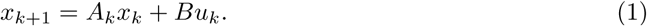

In this case, the three items listed above correspond respectively to the value of the state *x*_*k*_ at time *k*, the time dependent state transition matrix *A*_*k*_, and the input matrix *B* (along with the input function *u*_*k*_).

## Methods

### **Genome State Representation and Dimension Reduction: x***_k_*

The state representation *x* in Eq. 1 is the foundation for any control system and is critical for controllability analysis. To fully represent the state of a cell, a high number of measurements would need to be taken, including gene expression, protein level, chromatin conformation, and epigenetic measurements. As an initial simplification, we assume that the gene expression profile is a sufficient representation of the cell state.

Gene expression for a given cell is dependent on a number of factors, including (but not limited to): cell type, cell cycle stage, circadian rhythm stage, and growth conditions. In order to best capture the natural fibroblast dynamics from population-level data, time series RNA-seq was performed on cells that were cell cycle and circadian rhythm synchronized in normal growth medium conditions (See SI). Prior to data collection, all cells were temporarily held in the first stage of the cell cycle, G_0_/G_1_, via serum starvation. Upon release into the cell cycle, the population was observed every Δ*t* =8 hours (h) for 56 h, yielding 8 time points (at 0, 8, 16,…, 56 h). Let *g*_*i,k*_ be the measured activity of gene *i* = 1, *…, N* at measurement time *k* = 1, *…,* 8, where *N* is the total number of human genes observed (22,083). Analysis of cell-cycle marker genes indicated that the synchronized fibroblasts took between 32-40 h to complete one cell cycle post growth medium introduction, after which cells became largely unsynchronized (Fig. S1). Because of this, we define *K* =5 to be the total number of time points used for this model.

An obstacle to using *g* to represent *x* in a dynamical systems approach is the computational feasibility of studying a system with over 20,000 variables, necessitating a dimension reduction. Naïve dimension reductions such as partitioning the genome into 1 mega-base pair (Mb) bins ignores inherent structural organization of the genome and obscures important intricacies of finer resolutions. A comprehensive genome state representation should include aspects of both structure and function, and simultaneously have low enough dimension to be computationally reasonable. Along these lines, we propose a biologically inspired dimension reduction based on topologically associated domains (TADs).

The advent of genome-wide chromosome conformation capture (Hi-C) allowed for the studying of higher order chromatin structure and the subsequent discovery of TADs [8]. TADs are inherent structural units of chromosomes: contiguous segments of the 1-D genome for which empirical physical interactions can be observed [9]. Moreover, genes within a TAD tend to exhibit similar activity, and TAD boundaries have been found to be largely cell-type invariant [9, 10, 11]. TADs group structurally and functionally similar genes, serving as a natural dimension reduction that preserves important genomic properties. Fig. 1 depicts an overview of this concept. We computed TAD boundaries from Hi-C data via an algorithm that uses Fielder vector partitioning, described in Chen *et al.* (See SI) [12].

**Figure 1:**
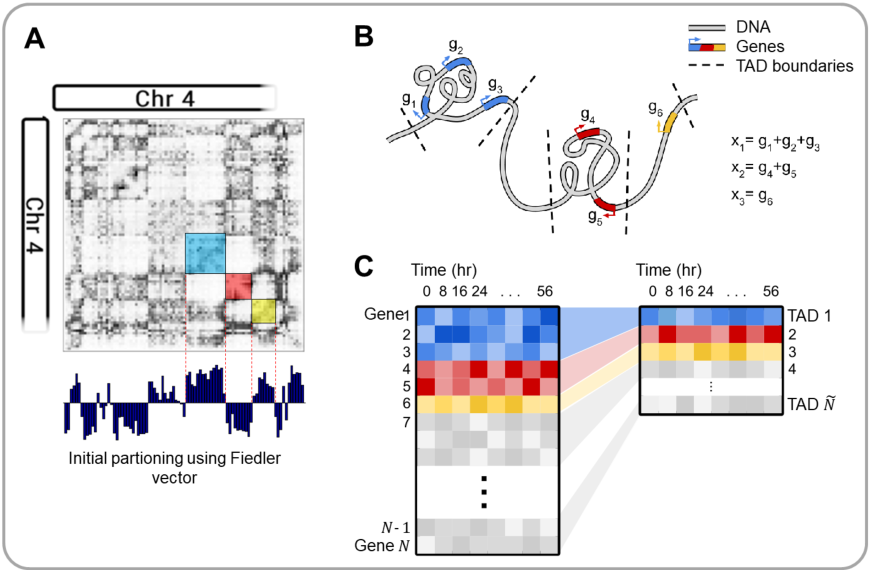
Overview of TAD dimension reduction. (*A*) Partitioning the Hi-C matrix based on the Fiedler vector. (*B*) Cartoon depiction of TAD genomic structure. (*C*) TAD dimension reduction summary.

Let *tad*(*i*) := *j* if gene *i* is contained within TAD *j*. We define each state variable *x*_*j,k*_ to be the expression level of TAD *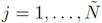* at time *k*, where *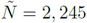* is the total number of TADs that contain genes. Specifically, *x*_*j,k*_ is defined as the sum of the expression levels of all genes within the TAD, measured in reads per kilobase of transcript per million (RPKM), i.e.

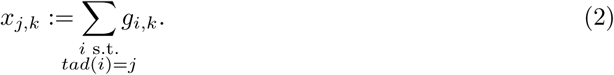

The vector of all TAD activities at measurement *k* is denoted with a single subscript *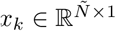*, *k* = 1,…, *K*.

### **State Transition Matrix: A***_k_*

Given the data we have, perhaps the most direct way to model the evolution of TAD activity level would be to assume a model of the form *x*_*k*+1_ = *x*_*k*_ + *y*, where *x*_*k*_ and *x*_*k*+1_ come from data, and *y* is the vector difference of *x*_*k*+1_ and *x*_*k*_. However, the data could also be viewed in a different way. Taken over a full cycle, the average value of the expression level of the 2,245 TADs is known, within experimental error. Assuming that there is a function *f* which maps *x*_*k*_ to *x*_*k+1*_, we can subtract the steady state average, 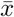, and focus on measuring the deviation from average as the cycle evolves. With this in mind, we have *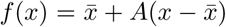* where *A* is allowed to depend on *x*’s location in the cell cycle. That is, we build a model for the variation of the cell cycle about *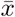*. For the model to match data and capture variability over the cell cycle, we will need to have a different *A* for each time step. Using the principle that *A* should differ as little from the identity as possible, we let *A*_*k*_ be the identity plus a rank one matrix chosen to match the data, for each time step *k*. In this case we have *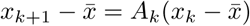*

We define a time dependent state transition matrix *A*_*k*_.

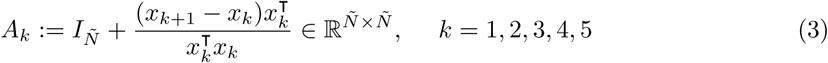

where *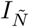* is the *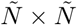* identity matrix.

### **Input Matrix and Input Signal: B**, **u***_k_*

With the natural TAD-level dynamics established in the context of our control Eq. 1, we turn our attention to quantifying methods for control.

A TF is a protein that can regulate a gene positively or negatively by binding to a specific DNA sequence near a gene and encouraging or discouraging transcription. This is accomplished, for example, by altering local chromatin conformation or by recruiting RNA polymerase II and other transcriptional machinery [13]. The degree to which a TF activates or represses a gene depends on the specific TF-gene interaction and most likely on a variety of nuclear subtleties and| | " intricacies that are difficult to quantify. Let *w*_*i,m*_ be the theoretical *regulation weight* of TF *m* on gene *i*, where *w*_*i,m*_ *>* 0 (*w*_*i,m*_ *<* 0) if TF *m* activates (represses) gene *i*, and *m* = 1, *…, M*, where *M* is the total number of well-characterized TFs. Weights that are bigger in absolute value, |*w*_*i,m*_ | *≫*0, indicate stronger transcriptional influence, and weights equal to zero, *w*_*i,m*_ = 0, indicate no influence.

Extensive TF perturbation experiments would be needed to determine *w*_*i,m*_ for each TF *m* on each gene *i*. Instead, we propose an alternative (simplified) method to approximate *w*_*i,m*_ from existing, publicly available data for TF binding sites, gene accessibility, and average activator/repressor + activity. To determine the number of possible binding sites a TF *m* recognizes near gene *i*, we scanned the reference genome for the locations of potential TF binding sites (TFBSs) (See SI).±Position frequency matrices (PFMs), which give information on TF-DNA binding probability, were obtained for 547 TFs from a number of publicly available sources (∴ *M=* 547). Let *c*_*i,m*_ be the number of TF *m* TFBSs found within ± 5kb of the transcriptional start site (TSS) of gene *i* (Fig.S2). In our model, the magnitude of *w*_*i,m*_ is proportional to *c*_*i,m*_. False negatives would include distal TFBSs outside of the ±5kb window, while false positives would be erroneous TFBS matches.

Although many TFs can do both in the right circumstances, most TFs have tendency toward either activator or repressor activity [13]. That is, if TF *m* is known to activate (repress) most genes, we can say with some confidence that TF *m* is an activator (repressor), so *w*_*i,m*_ *≥*0 (*w*_*i,m*_ *≤*0) for all *i*. To determine a TF’s function, we performed a literature search for all 547 TFs and labeled 299 as activators and 124 as repressors (See SI). The remaining TFs were labeled unknown for lack of conclusive evidence for activator or repressor function. In the case of inconclusive evidence, the TF was evaluated as both an activator and a repressor in separate calculations. Here, we define *a*_*m*_ as the activity of TF *m*, with 1 and -1 denoting activator and repressor, respectively, and the sign of *w*_*i,m*_ will be determined by *a*_*m*_.

TFBSs are cell-type invariant since they are based strictly on the linear genome. However, it is known that for a given cell type, certain areas of the genome may be opened or closed depending on epigenetic aspects. To capture cell type specific regulatory information, we obtained gene accessibility data through DNase-seq. DNase-seq extracts cell type specific chromatin accessibility information genome-wide by testing the genome’s sensitivity to the endonuclease DNase I, and sequencing the non-digested genome fragments. This data is used for our initial cell type to determine which genes are available to be controlled by TFs [14]. Here, we define *s*_*i*_ be the DNase I sensitivity information (accessibility; open/close) of gene *i* in the initial state, with 1 and 0 denoting accessible and inaccessible, respectively (See SI).

We approximate *w*_*i,m*_ as

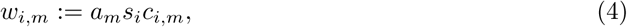

so that the magnitude of influence is equal to the number of observed consensus motifs *c*_*i,m*_, except when the gene is inaccessible (*s*_*i*_ = 0) in which case *w*_*i,m*_ = 0.

Since we are working off a TAD-dimensional model, our input matrix *B* must match this dimension. Let *b*_*m*_ be a 2,245-dimensional vector, where the *j*^*th*^ component is

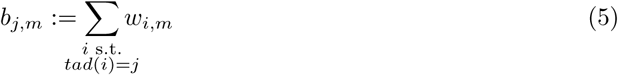

and define a matrix *B* = [*b*_1_ *b*_2_… *b*_*M*_].

The amount of control input is captured in *u*_*k*_, which is a ℝ^*M×*1^ vector representing the quantity of the external TFs we are inputting to the system (cell) at time *k*. This can be controlled by the researcher experimentally through manipulation of the TF concentration [15]. In this light, we restrict our analysis to *u*_*k*_*≥* 0 for all *k*, as TFs cannot be subtracted from the cell. *u*_*m,k*_ is defined as the amount of TF *m* to be added at time point *k*.

With all variables of our control Eq. 1 defined, we can now attempt to predict which TFs will most efficiently achieve cellular reprogramming from some *x*_*I*_ (initial state; fibroblast in our setting) to *x*_*T*_ (target state; any human cell type for which compatible RNA-seq data is available) through manipulation of *u*_*k*_. An overview of our DGC framework is given in Fig. 2.

**Figure 2:**
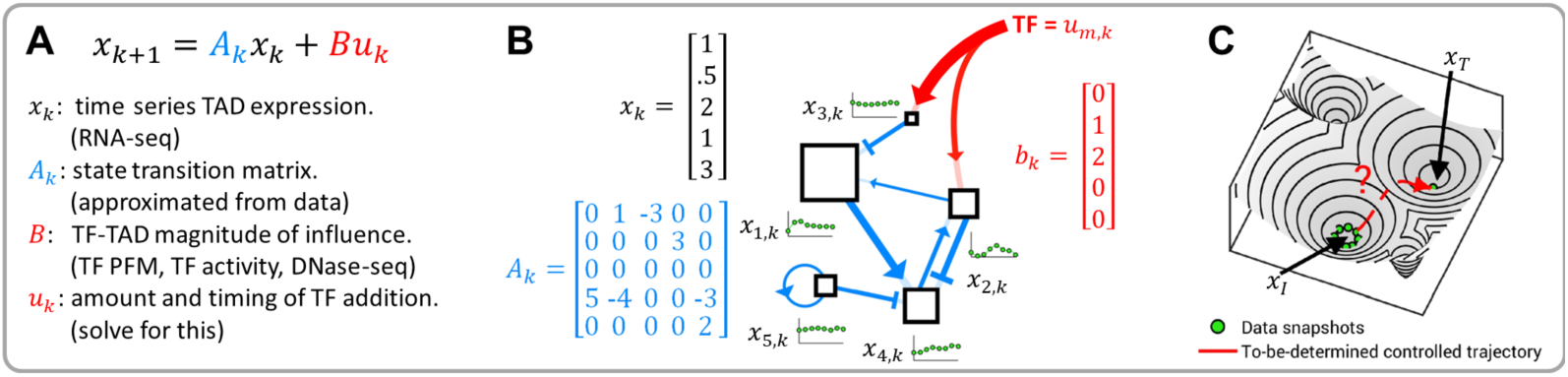
Data-guided control overview. (*A*) Summary of control equation variables. (*B*) Each TAD is a node in a dynamic network. The blue connections represent the edges of the network and are determined from time series fibroblast RNA-seq data. The miniature green plots represent the expression of each TAD changing over time. The red arrows indicate additional regulation imposed by exogenous transcription factors. (*C*) A conceptual illustration of the problem: can we determine transcription factors to push the cell state from one basin to anotherffi

### Selection of TFs

We consider different scenarios for the type of input regime. The first is assuming the input signal is constant *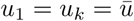* intended to mimic empirical regimes where TFs are given at a single time point. We will show that this method is theoretically inferior to inputting different TFs at different points in the cell cycle in a later section.

Eq. 1 has an explicit solution that is easily computed.

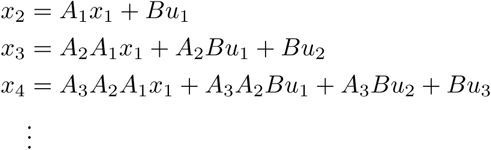

Notice the expression for *x*_4_ depends the input matrix *B* and the input signal *u*_*k*_.

If *x*_*T*_ is a target condition, then the Euclidean distance ║ ║ can be used to measure how close a state is to the target state, i.e.:

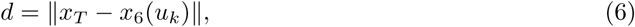

where the notation *x*_6_(*u*_*k*_) is used to emphasize the dependence of *x*_6_ on *u*_*k*_. Considering all possible input signals, one can compute the optimal control that finds the minimum distance for a given initial and target cell type. Let *u*_*\*k*_ denote the optimal *u*_*k*_ used to minimize *d*, and *d*_*_ denotes this minimum distance value.

We note here that when determining which TFs can be used to reach *x*_*T*_, it is often desirable and more experimentally feasible to minimize the number of distinct TFs given to the cells. Transfection of cells with multiple different TFs can lead to cell stress and death, and a lower efficiency of transfection overall. Moreover, many experimentally confirmed direct reprogramming regimes use ≤ 4 TFs to achieve reprogramming. For these reasons, we set all indices of *u*_*k*_ equal to zero, except for indices corresponding to TFs that we choose.

We define *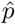* to be a set of positive integers that refer to the indices of *u*_*k*_ (read: TFs) that are allowed to be non-zero (e.g. *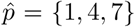* = {1, 4, 7} refers to TFs 1, 4, and 7). Let *p* be the number of elements in *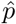*. Given a set of TFs *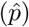*, we determine the quantity and timing of TF input (*u*k*) that minimizes the difference (*d\**) between the initial (*x*_*I*_) and target (*x*_*T*_) cell state. Mathematically, this can be written as

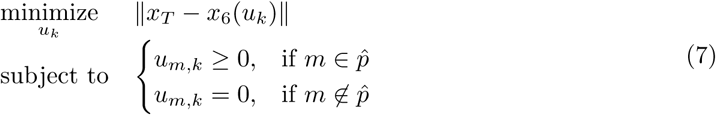

We use MATLAB’s *lsqnonneg* function to solve Eq. 7, which gives *u*k* and *d\**.

Let *d*_0_ := ∥*x*_*T*_– *x*_*I*_ ∥, be the distance between initial and target states with no external influence. We define the score *μ* := *d*_0_ *d\**, which can be interpreted as the *distance progressed towards target μ* can be calculated for each *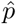* and sorted (high to low) to determine which TF or TF combination is the best candidate for direct reprogramming between *x*_*I*_ and *x*_*T*_.

**Remark:** Subsets of TFs were chosen for each calculation based on the following criteria:. =10-fold expression increase in target state compared to initial state, and. =4 RPKM in target state. These criteria are used to select differentially expressed TFs and TFs that are sufficiently active in the target state.

## Results

### Quantitative Measure Between Cell Types

In order to best utilize our algorithm to predict TFs for reprogramming, compatible data on target cell types must be collected. For this, we explore a number of publicly available databases where RNA-seq has been collected, along with RNA-seq data collected in our lab. The ENCODE Consortium has provided data on myotubes and embryonic stem cells (ESCs) (See SI) [16]. The GTEx portal provides RNA-seq data on a large variety of different human tissue types [17]. Although each GTEx experiment is performed on tissue samples, thus containing multiple different cell types, we use these data as more general cell state targets.

To give a numerical structure to cell type differences, conceptually similar to Waddington’s epigenetic landscape, we calculate *d*_0_ between all cell types collected. Fig. 3*A* shows *d*_0_ values for 32 tissue samples collected from the GTEx portal, along with ESC, myotube, and our fibroblast data (additional cell type *d*_0_ values shown in SI). Warmer colors (red) denote further distances between cell types. GTEx RNA-seq data is scaled to keep total RPKM difference between time series fibroblast and GTEx fibroblast RNA-seq minimal (See SI).

**Figure 3:**
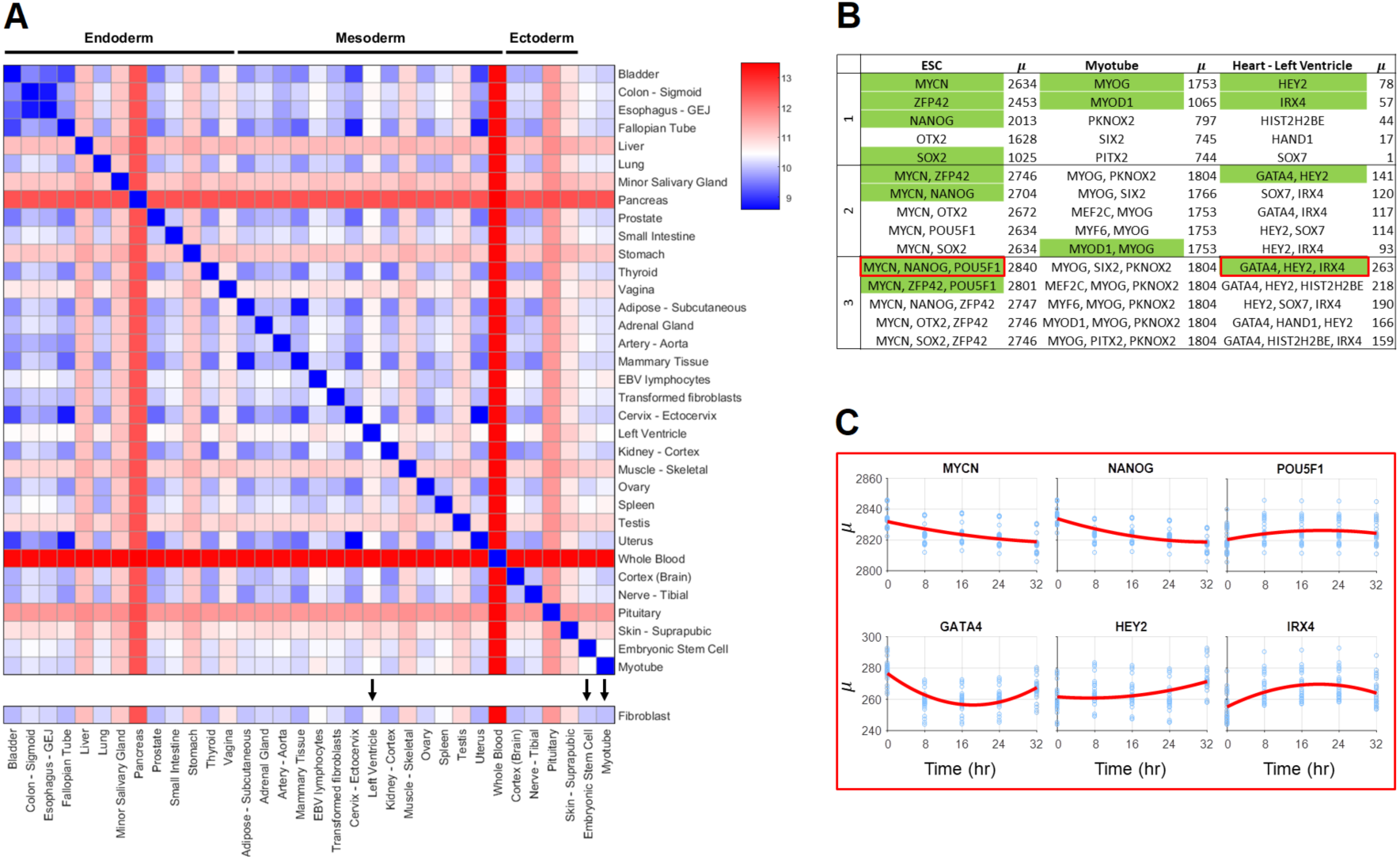
Quantitative measure between cell types and transcription factor scores. (*A*) *d*_0_ values between GTEx tissue types and ESC, myotube, and fibroblast. Tissue types and cell types with black arrows have predicted transcription factors for reprogramming from fibroblasts shown in 3B. (*B*) Table of predicted transcription factors for a subset of cell and tissue types. Top 5 transcription factors for combinations of 1-3 shown. Green labeled transcription factors are either highly associated with the differentiation process of the target cell type and/or validated for reprogramming. These transcription factors are discussed in the main text. (*C*) Time-dependent scores for selected combinations of 3 transcription factors for fibroblast to ESC and fibroblast to “Heart Left ventricle". x-axis refers to time of transcription factor addition, y-axis refers to *μ*.

### TF Scores

To assess our method’s predictive power, a subset of target cell types are presented here that have either validated TF reprogramming methods or TFs highly associated with the target cell type. Additional predicted TFs for reprogramming are included in SI. We note that though experimentally validated TFs provide the best current standard for comparison, we believe experimental validation with our predicted TFs may provide more efficient reprogramming results. For all reprogramming regimes presented in this section, fibroblast is used as the initial cell type due to the availability of synchronized time series data, and all TFs are introduced at *k* =1 [9].

For conversion of fibroblast to myotubes, the top predicted single input TFs are MYOG and MYOD1, both of which are known to be crucial for myogenesis. While MYOD1 is the classic master regulator reprogramming TF for myotube conversion, activation of downstream factor MYOG is necessary for full conversion [18]. For fibroblast to ESC conversion, a number of TFs known to be necessary for pluripotency are predicted, including MYCN, ZFP42, NANOG, and SOX2 [19]. With the knowledge that no single TF has been shown to fully reprogram a fibroblast to an embryonic state, combinations of TFs are more informative for this analysis. The top scoring combination of 3 TFs is MYCN, NANOG, and POU5F1–three well-known markers for pluripotency [19]. Interestingly, POU5F1 scores poorly when input individually, but is within the top set of 3 TFs when used in combination with MYCN and NANOG. Left ventricle reprogramming includes TFs that are known to be necessary for natural differentiation in the top score for all 1-3 combinations. These include GATA4 (a known TF in fibroblast to cardiomyocyte reprogramming), HEY2, and IRX4 [20, 21, 22].

### Time-dependent TF Addition

Fibroblast to ESC conversion was of particular interest in our analysis as this is a well-studied regime with a number of validated TFs (with a variety of reported efficiencies), and this conversion is promising for its regenerative medicine application. High scoring TFs yield many that are known markers for pluripotency, but the top combination of 3, MYCN, NANOG, and POU5F1, has not been used specifically together, to our knowledge.

Since our method incorporates dynamical RNA-seq data, analysis can be extended to determine the best time to input control for a given set of TFs. In our model, there are five possible input times: 0, 8, *…*, 32 h. We assume a TF continues to influence the system at a constant value once input until the final time (40 h). We restrict our analysis here to combinations of 3 TFs. This gives 5^3^ = 125 possible *Time-dependent regimes* to input the TFs; e.g. *TF1, TF2, TF3* are input, respectively, at times 0,0,0, or 0,0,8, or 0,0,16, or *…* or 32,32,24 or 32,32,32. Inputting a TF at time *k*^*^ can be viewed mathematically as requiring *um,k* =0 for all *k* < *k*^*^.

Time-dependent analysis of the top scoring ESC TFs reveals that scores vary widely depending on the time of input. MYCN and NANOG show a strong preference for input at the beginning of the cell cycle, while POU5F1 shows a slight preference for input towards the end of the cell cycle, with the highest score achieved when MYCN and NANOG are input at 0 h and POU5F1 is input at 32 h. Analysis on how the time of input control affects *μ* is shown in Fig. 3*C*. Time-dependent analysis was also conducted for the top combination of 3 TFs for fibroblast to left ventricle. This set includes GATA4, HEY2, and IRX4, all factors highly associated with the cardiac phenotype [20, 21, 22]. This analysis predicted that the best reprogramming results would occur if GATA4 is given immediately (0 h), with IRX4 and HEY2 given later (24 and 32 h, respectively).

## Discussion

The results from this algorithm show promise in their prediction of known reprogramming TFs, and demonstrate the importance of including time series data for gene network dynamics. Time of input control has shown to have an impact on the end cell state, in line with what has been shown in natural differentiation [23].

While we believe that this is the best model currently available for predicting TFs for reprogramming, we are aware of its limitations and assumptions. TAD-based dimension reduction is based on the observation that genes within them correlate in expression over time, though we lack definitive proof of regulation by shared transcriptional machinery [9]. This assumption was deemed necessary for dimension reduction in the context of deriving transition matrix *A*_*k*_. With finer time steps in RNA-seq data, the assumption may not be necessary for TF prediction, at the cost of increased computation time. Additionally, a 5kb window flanking the TSS of each gene was used to ensure that all potential regulators are found, at the cost of potential inclusion of false positive motifs.

GTEx data proved to be an invaluable resource for testing our algorithm, providing many target states for TF prediction. It is important to note that these data are collected from cadaver tissue samples; therefore the RNA-seq data is coming from a heterogeneous cell-type population and may be enriched for stress factors known to be elevated after death (e.g. HSF4). Ideally, RNA-seq data for target cells would be derived from a homogeneous population, with minimal experiment collection variables. Future work includes the extension of this DGC approach to other target cell types.

Although this program can score TFs relative to other TFs in a given reprogramming regime, it is difficult to predict a *μ* threshold that would guarantee conversion. Additionally, rigorous experimental testing will be required to validate these findings and determine how our *u* vector translates to TF concentration. This is a product of the large number of assumptions that must be made to develop the initial framework for a reprogramming algorithm. With finer resolution in the time series gene expression, more subtle aspects of the genomic network may be observed, allowing for better prediction.

Our proposed data-guided control framework successfully identified known TFs for fibroblast to ESC and fibroblast to muscle cell reprogramming regimes. The framework rates individual TFs as well as sets of TFs. We employ a biologically-inspired dimension reduction via TADs, a natural partitioning of the genome. This comprehensive state representation was the foundation of our framework, and the success of our methods motivates further investigation of the importance of TADs as functional units to control the genome.

A dynamical systems view of the genome allows for analysis of timing, efficiency, and optimality in the context of reprogramming. Our framework is the first step toward this view. The successful implementation of time-varying reprogramming regimes would open new avenues for direct reprogramming. Experimental verification of predicted regimes and development of methods to identify optimal sets of TFs are planned for the near future. This template can be used to develop regimes for changing any cell into any other cell, for applications that include reprogramming cancer cells and controlling the immune system. Our DGC framework is well equipped for designing personalized cellular reprogramming regimes. Finally, this framework can serve as a general technique for investigating the controllability of networks strictly from data.

## Methods and Materials

Hi-C and RNA-seq data were collected from cell cycle and circadian rhythm-synchronized proliferating human fibroblasts of normal karyotype. Data were collected every 8 h, spanning 56 h. Publicly available data was used for target cell types. Detailed materials and methods are provided in Chen *et al.* and in SI [9].

## Acknowledgement

We thank Robert Oakes, Emily Crossette, and Sijia Liu for their critical reading of the manuscript and helpful discussions. We extend special thanks to James Gimlett and Srikanta Kumar at Defense Advanced Research Projects Agency (DARPA) for support and encouragement. This work is supported, in part, by the DARPA Biochronicity Program and the DARPA Deep-Purple and FunCC Program.

